# Label-free single-cell live imaging reveals fast metabolic switch in T lymphocytes

**DOI:** 10.1101/2023.01.04.522685

**Authors:** Noémie Paillon, Thi Phuong Lien Ung, Stéphanie Dogniaux, Chiara Stringari, Claire Hivroz

## Abstract

T cell activation induces a metabolic switch generating energy required for proliferation, survival, and fueling their functions. Thus, it is essential to monitor metabolism associated to subcellular functional and structural changes. We used non-invasive label-free two-photon fluorescence lifetime microscopy (2P-FLIM) to map the spatial and temporal dynamics of the metabolic NADH co-enzyme during T lymphocyte activation. 2P-FLIM measurements of the protein-bound and free NADH ratios provides a readout of the redox state (NAD^+^/ NADH) of the cells, and thus of their OXPHOS and glycolysis rates. Using this method, we followed the dynamics of fraction of bound NADH (fb NADH) in live single cells. Comparing fb NADH between resting and activated T cells, we show that T cell activation induces a rapid switch toward glycolysis. The switch takes only 10 minutes and remains stable for at least one hour. Three-dimensional (3D) analysis revealed that the intracellular distribution of fb NADH is symmetrically distributed in resting cells, whereas increases at the contact zone in activated cells. Finally, we show that fb NADH negatively correlates with spreading of activated T cells, suggesting a link between actin remodeling and metabolic changes. This study shows that 2P-FLIM measurement of fb NADH is well suited to follow a fast metabolic switch in 3D, in single T lymphocytes with subcellular resolution.

## INTRODUCTION

T lymphocyte activation through the T cell receptor (TCR) and co-receptors such as CD28 induces a rapid transcription of numerous new messenger RNA (mRNA) transcripts and proteins. Activated T lymphocytes also undergo massive growth, doubling to quadrupling their size in 1 to 2 days as well as several cycles of divisions^1^. These different steps are demanding in energy and thus require a rewiring of T lymphocyte metabolism^2^. Indeed, T lymphocytes change their metabolism during immune response. Briefly naïve T cells that are metabolically quiescent, mainly rely on oxidative phosphorylation (OXPHOS) for their energetic needs. Effector T cells, which are very metabolically active have higher rates of both glycolysis and OXPHOS, whereas memory T cells, which are quiescent but metabolically primed rely on fatty acid oxidation and OXPHOS^2^. The field of immunometabolism has gained importance by showing that different T lymphocyte populations have different metabolic signatures, but also by unveiling that the metabolism does not only energetically supports cellular functions but also shape them^3^. These better understanding of immunometabolism of T cells paved the way to harnessing metabolism for therapeutic interventions. For examples, finding the right conditions to generate CAR T cells that are “metabolically fit”^4^ or overcoming metabolic competition in the tumors microenvironment to improve anti-tumoral T cell responses^5^. There is thus an increasing need to better understand immunometabolism in T lymphocytes.

Many tools have been developed to measure cell metabolism and have been used in T lymphocytes. Among them, we can cite 13 C-based stable isotope labeling (SIL) techniques that facilitate tracing the metabolic fate of nutrients in cells^6^, the Seahorse Extracellular Flux Analyzer, which simultaneously measures, in real-time, the extracellular acidification rate (ECAR; an indicator of glycolysis) and the oxygen consumption rate (OCR; an indicator of oxidative phosphorylation (OXPHOS)) from relatively low numbers of cells. Both these methods allow the metabolic analysis of cells in bulk. We can also cite the two recent elegant assays: SCENITH and SPICE-Met that rely respectively on – characterizing the energetic metabolism profile of cells by monitoring changes in protein synthesis levels in response to metabolic inhibitors^7^, or on – sensing the ATP:ADP ratio in cells expressing the genetically encoded fluorescent biosensor^8^. The SCENITH assay, based on flow cytometry, monitors the metabolic profile at a single cell resolution but not in real time and has been used successfully in T cells^9^. The SPICE-Met assay is based on one or two-photon microscopy, can be used in real-time imaging on single cells, has been used successfully in T cells^10^ but requires the expression of a transgene.

Most of the studies aiming at analyzing metabolism in T lymphocytes have been performed in cell populations at steady state or several hours after exposure to activation signals. Rapid changes (in the order of 30 to 60 min) in OXPHOS and aerobic glycolysis have yet been reported upon activation of T lymphocytes ^11,12^.

To tackle both the heterogeneity of the metabolic profiles and of their temporal dynamic, we used Two-photon Fluorescence Lifetime Microscopy (2P-FLIM) of the coenzyme NADH, as it reports on metabolic processes in a label-free and non-invasive way^13–15^. NADH is the principal electron acceptor in glycolysis in oxidative phosphorylation and its ubiquity renders NADH one of the most useful and informative intrinsic biomarkers for metabolism in live cells and tissues^16,17^. Through the readout of protein-bound and free NADH, 2P-FLIM provides sensitive measurements of the redox states (NAD^+^/ NADH) of cells as well as OXPHOS and glycolysis rates^14,15,18–22^. 2P-FLIM has the spatio–temporal resolution required to characterize NADH subcellular compartments^23^ and to measure metabolic shifts induced upon T lymphocyte activation^24^ as well as metabolic polarization and heterogeneity in macrophages and microglia *in vitro*^25–27^ as well as *in vivo* ^28–31^.

In the present study, we used 2P-FLIM of NADH to follow the dynamic of NAD^+^/ NADH redox ratio (through the fraction of bound NADH) in Jurkat T cells (a leukemic T cell line) and primary human CD4^+^ T lymphocytes, during activation. We activated the cells on glass slides coated with anti-CD3 + anti-CD28 antibodies, which induce polarization and spreading of T cells and mimic the immune synapse formation^32^. Our results show that T cell activation induces a rapid decrease of fraction of bound NADH, reflecting a decrease in oxidative phosphorylation/glycolysis ratio in activated T lymphocytes. This can be followed, upon time, at a single cell level and in 3D. The three-dimensional metabolic maps show that the distribution of fraction of bound NADH in the cell is changing in different cellular compartments upon immune synapse formation. Moreover, because remodeling of the actin cytoskeleton is one of the early hallmarks of T cell activation we expressed LifeAct-mCherry, a 17-amino-acid peptide, which stained filamentous actin (F-actin) structures in Jurkat T cells and followed the bound/free NADH ratio simultaneously with the spreading. This analysis showed that the fb NADH diminished in spreading cells suggesting that there might be a link between actin remodeling in T cells and metabolic changes.

Together, our data show that upon activation mimicking immune synapse formation, T lymphocytes dynamically modify their metabolism in time and space.

## MATERIALS AND METHODS

### Cells

Jurkat T cells (94% homology with Jurkat clone E6.1 validated by the SSTR method on the DSMZ website) were cultured at 37°C with 5% CO_2_ in RPMI 1640 Glutamax (Gibco; 61870-010) supplemented with 10% fetal calf serum (Eurobio; CVFSVF00-01) and were passed every 2-3 days at approximatively 0.5 × 10^6^ cells/mL.

Peripheral blood mononuclear cells (PBMCs) from healthy donors were isolated using a ficoll density gradient. Buffy coats from healthy donors were obtained from Établissement Français du Sang in accordance with INSERM ethical guidelines. CD4^+^ T cells were purified using the total CD4^+^ negative isolation kit (Miltenyi Biotech; 130-096-533). Primary CD4^+^ T cells were activated in six-well plates coated with anti-CD3e (OKT3; 317326; BioLegend) in presence of soluble anti-CD28 (CD28.2; 302914; BioLegend) during 6 days prior to use and cultured in RPMI 1640 Glutamax (Fisher Scientific; 61870-010) supplemented with 10% fetal calf serum (Eurobio, CVFSVF00-01), 10 000 U/mL penicillin-streptomycin (Gibco; 15140-122), 1M Hepes (Gibco, 15630-056), 50mM 2-Mercaptoethanol (Gibco; 31350-010) and 20 U/mL recombinant IL-2 (Immunotools; 11340025).

### Lentivirus production and Jurkat cell transduction

Replication-defective lentiviral particles were obtained by transfecting HEK-293T cells with Gag, Pol, rev, encoding plasmid (pPAX2), envelop encoding plasmid (pMD2.G) and the LifeAct-mCherry construct encoded in a pWXLD vector. After 48h, lentiviruses were recovered in the supernatant and concentrated. Jurkat cells were transduced and the positive fraction was sorted by flow cytometry (SH800 Cell Sorter, Sony Biotechnology) to establish a stable LifeAct-mCherry^+^ Jurkat cell line.

### T cell activation

Poly-d-lysine coated 35mm glass-bottom dishes (MatTek; P35GC-1.0-14-C) were coated with 0,1μg/mL anti-CD3 (OKT3; 317326; BioLegend) and 10μg/mL anti-CD28 (CD28.2; 302914; BioLegend) at 4°C overnight and washed with PBS before use. 300 000 cells were resuspended in RPMI 1640 without phenol red (Fisher Scientific, 10363083) supplemented with 10% fetal calf serum and plated on coated dishes prior imaging.

### Two-photon excited fluorescence lifetime imaging (FLIM)

Imaging was performed on a laser scanning microscope (TriMScope, Lavision Biotec, Germany). A simplified scheme of the multiphoton microscope is shown in Fig. 1A. The excitation is provided by a dual-output femtosecond laser (Insight DS++, Spectra-Physics, Santa Clara, CA, USA) one with a first beam tuneable from 680 nm to 1300 nm (120 fs pulses, 80 MHz) and a second one with a fixed wavelength beam at 1040 nm (200 fs pulses). A water Immersion objective (25X, NA=1.05, <u>XLPLN-MP</u>, Olympus, Japan) is used to focus the laser on the sample and collect fluorescence signal. Fluorescence signal is epi-detected by a hybrid photomultiplier tube (R10467U, Hamamatsu, Japan).

**Fig. 1:**
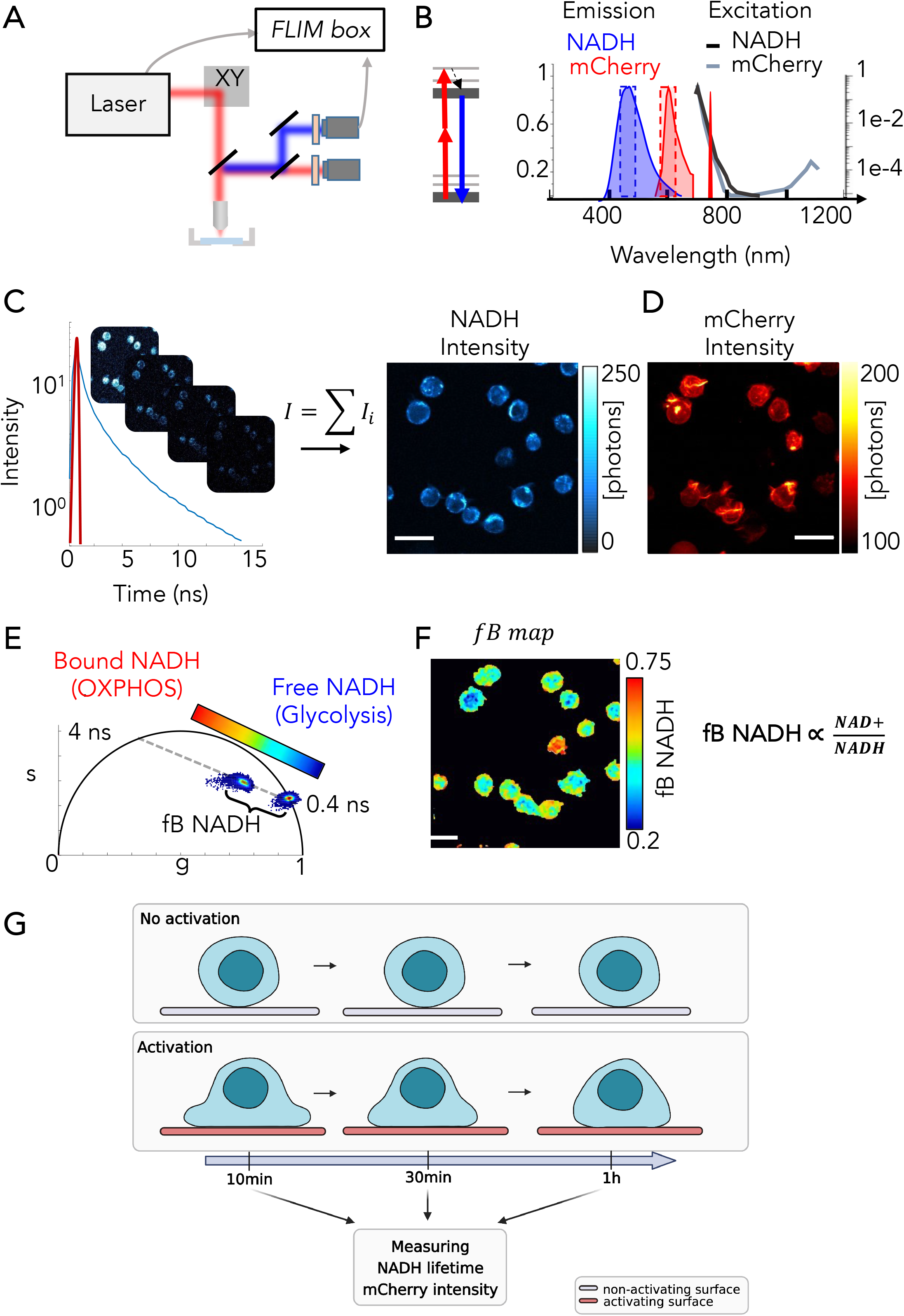
Experimental setup of *two-photon excitation Fluorescence Lifetime Imaging (FLIM)*. **A**. Scheme of the experimental setup used for this work. **B**. Principles of two-photon simultaneous excitation of NADH and LifeAct-mCherry fluorophores at 760 nm. Emission fluorescence of NADH and LifeAct-mCherry collected with band-pass filters 460/80 and 607/70 respectively. **C**. Fluorescence lifetime image contains a fluorescence lifetime decay in every pixel. The NADH intensity image is determined by the sum of the FLIM stack. **D**. LifeAct-mCherry intensity image simultaneously acquired in the red channel. **E**. The multi-exponential fluorescence intensity decay in every pixel of the image is transformed in the phasor plot by Fourier transform. Calculation of the fraction of bound NADH (fb NADH) is performed by measuring the distance of every pixel from the location of free NADH. **F**. Fraction of bound NADH is proportional to the redox ratio NADH/NAD+ ratio. **G**. Schematic representation of T-cell activation.

To perform fluorescence lifetime microscopy of NADH, 740nm wavelength excitation was used with a typical power of 12 mW. A blue band-pass filter was installed in front of the detector to collect NADH autofluorescence (Semrock FF01–460/80). The red fluorescence of LifeAct-mCherry is simultaneously collected with a red band pass filter (Semrock FF01– 607/70). Time-correlated single photon counting (TCSPC) electronics (Lavision Biotec, Germany) measures the arrival time of the fluorescence photons with respect to the laser pulse and reconstructs the fluorescence lifetime decay. The laser trigger is taken from the fixed wavelength beam using a photodiode (PDA10CF-EC, Thorlab). Lifetime calibration of the FLIM system was performed by measuring the lifetime of fluorescein at pH9 with a single exponential of 4.04 ns (Fig.1C). We typically collected 200-300 photons during an acquisition time in the order of one minute for a 256×256 pixels image at a pixel dwell time of 60.8 μs/pixel. The three-dimensional (3D) FLIM imaging was performed with a Z-step of 3 μm and the total acquisition time of a 3D FLIM is in the order of two minutes.

### T Lymphocytes imaging

T lymphocytes were imaged on a glass-bottom dish, containing a non-fluorescent assay medium (Fisher Scientific, 10363083) placed inside an incubation chamber at 37 °C and 5% CO_2_ (Okolab, Pozzuoli, Italy).

We performed label-free fluorescence lifetime microscopy of NADH in a total of 87 ROIs with 700 cells in 2 independent experiments for the stable cell line and in a total of 56 ROIs with 1000 cells in 1 independent experiment for the primary cells.

To measure the metabolic trajectory in cells we treated the cells with a solution of 50 μM of rotenone (R8875; Sigma-Aldrich, St. Louis, MO, USA) to block the respiratory chain via complex I and 2-deoxy-D-glucose (D8375-10MG; Sigma-Aldrich, St. Louis, MO, USA) of 10mM to inhibit glycolysis.

### Analysis of the Fluorescence Lifetime microscopy images

Intensity data were analyzed and treated with Fiji-ImageJ (NIH, Bethesda, MD, USA). All FLIM data were processed and analyzed with a custom written software Matlab (Mathworks, Natick, MA, USA). FLIM data were transformed from time domain to frequency domain by using a FFT and plotted in the phasor plot as previously described^14,33^. As our FLIM data were acquired in the time domain (Fig. 1C) we transformed the multi-exponential fluorescence intensity decay in every pixel of the image into a point in the 2D phasor plot of coordinates *g* and *s* (Fig.1E) with the following equations:

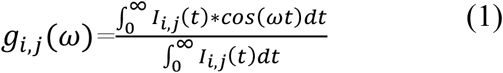

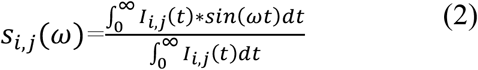

where i and j indicate the order pixel of the image and ω is the frequency. ω is calculated through the laser repetition rate *ω*=2*πf* (*f*=80 MHz). We calibrated our FLIM system by using a solution of fluorescein at pH 9 that has a single lifetime of 4.04 ns and we measured the lifetime of free NADH in solution of 0.4ns. The fraction of bound NADH is graphically calculated as the distance between the experimental point (*g*_*exp*_, *s*_*exp*_) and the location of free NADH (*g*_*fNADH*_, *s*_*fNADH*_) (Fig. 1E) using the following equation:

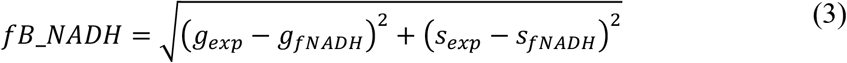

We performed 3D-FLIM single cell analysis using the red channel LifeAct-mCherry for the stable cell line and on the NADH intensity channel for the primary T cells (Fig. S1A). We first performed a Z projection of all the Z planes of the 3D stack (Fig. S1A). Using the LifeAct-mCherry channel, binary masks were created to identify and segment each single cell (Fig. S1B) using Fiji-ImageJ (NIH, Bethesda, MD, USA). Based on a Matlab custom written code, we applied the masks of single cells to the 3D fb NADH maps (Fig. S1C) to create 3D metabolic maps of single cells (Fig. S1D). We then calculated the average fb NADH for every Z plane of the cell and in the entire volume of the cell.

## RESULTS

### Two-photon fluorescence lifetime imaging (FLIM) of the metabolic coenzyme NADH in T lymphocytes

In this study, we implemented label-free two-photon fluorescence lifetime microscopy (2P-FLIM)^14,15^ in living T lymphocytes to create non-invasive metabolic maps of the metabolic coenzyme NADH at a submicron resolution and measured the spatial and temporal dynamics of redox metabolism during T lymphocyte activation.

The principle of 2P-FLIM NADH is described in Fig. 1. To obtain excitation of NADH and LifeAct-mCherry simultaneously, we excited the T lymphocytes in culture with an excitation wavelength of 760nm, while the emitted fluorescence was collected in the blue and red spectral ranges respectively (Fig. 1A, 1B and Material and Methods). Representative 2P NADH two-photon (2P) intensity images of live T lymphocytes and simultaneously acquired 2P LifeAct-mCherry intensity images are shown in Fig. 1C and Fig. 1D. We performed FLIM of NADH in T lymphocytes using a commercial time-correlated single photon counting (TCSPC) system and we established experimental conditions that allow non-invasive longitudinal metabolic imaging of the same cells without any photodamage (Fig. 1C and Materials and Methods). The fluorescence lifetime of NADH changes upon binding to enzymes within the electron transport chain. Indeed, the protein-bound NADH lifetime is significantly longer than the free NADH lifetime, due to self-quenching in the free state^13^. Thus, fluorescence lifetime imaging (FLIM) is used to provide sensitive measurements of the free and protein-bound NADH ratio to estimate the contribution of glycolysis versus OXPHOS in ATP production^15–17^.

The NADH FLIM images were analyzed through phasor analysis^14,34^(Fig. 1E). To quantify the subcellular metabolism, we calculated the fraction of bound NADH (fb NADH) in every pixel of the image (Fig. 1E and Material and Methods) with respect to free NADH, which is proportional to the redox ratio NAD+/NADH^19^ (see Material and Methods). The fb NADH map (Fig. 1F) highlights the metabolic heterogeneity among cells and the subcellular NADH compartmentalization. This experimental setup was applied to establish the cellular metabolic trajectory of human T lymphocytes in resting and activating conditions.

### FLIM imaging of the metabolic coenzymes NADH in resting and activated Jurkat T cells upon time

The aim of the study was to measure the temporal dynamics of fluorescence lifetime of NADH in T cells after activation, to determine the glycolytic capacity of T cells in real time. To do so, we used a model that has been developed to mimic activation of T cells at the immune synapse. Jurkat T cells, transduced with a lentiviral vector encoding LifeAct-mCherry to follow F-actin remodeling, were dropped on glass surfaces coated with either poly-L-Lysine (PLL) alone, or with PLL and anti-CD3 + anti-CD28 activating antibodies (Fig.1G). This activation is known to induce early signaling and is accompanied by spreading of T cells on the surface. Measurements were performed on Jurkat cells in 3D (Fig. S1 and Material and Methods) at 10, 30 and 60 minutes after plating the cells on PLL or activating antibodies (Fig. 1G). In the control, non-activated Jurkat cells plated on PLL, the distribution of the NADH two-photon fluorescence intensity presented a dim central area, corresponding to nuclei and a brighter cortical area reflecting the presence of mitochondria in the cytosol (Fig. 2A, upper raw). In fact, mitochondria have significantly higher NADH/NAD+ ratio (1:5 to 1:10) with respect to the nucleus and cytoplasm (1:400 to 1:700)^18,19^ and NADH is fluorescent while NAD+ is not. In the presence of anti-CD3+CD28 antibodies, the 2P fluorescence intensity of NADH revealed a more even distribution close to the activating slide, reflecting changes in the morphology of activated T cells with a possible accumulation of mitochondria at the contact zone (surrogate immune synapse).

**Fig. 2:**
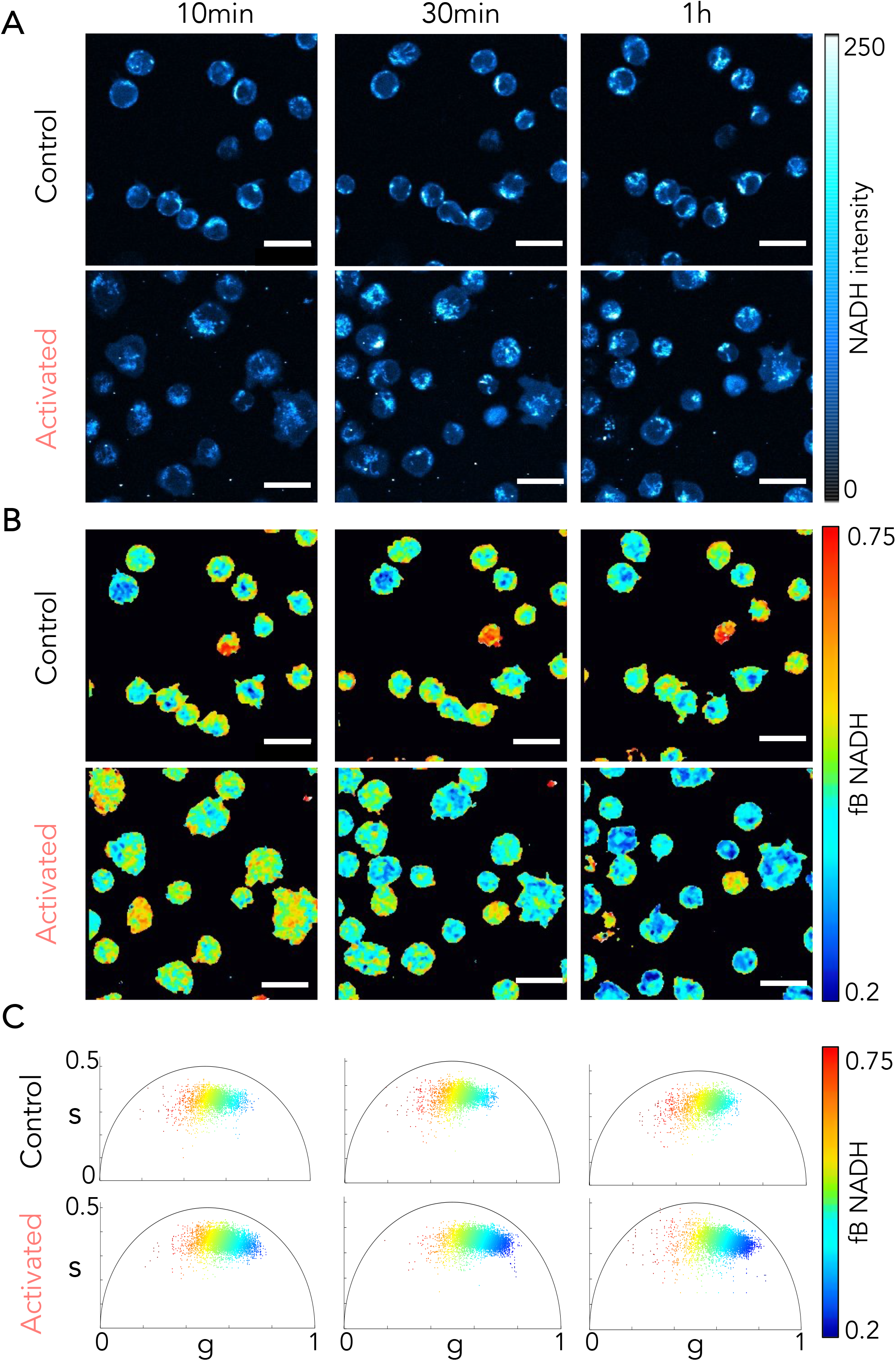
Metabolic shift during T cell activation revealed by NADH FLIM. **A**. Representative images of NADH intensity, **B**. maps of fraction of bound NADH and **C**. phasor plot scatters of T cells in control and activated condition at three time point (10 min, 30 min and 1 hour). Every single point in the phasor plot is color coded after the fraction of bound NADH.

The fb NADH maps were calculated in every Z plane of the cell (Fig. S1C) and the metabolic signature of the entire cell was evaluated by averaging the fb NADH in 3D (Material and Method). Whereas the fb NADH did not change over time in Jurkat cells plated on PLL (Fig. 2B, upper panel), there was a general decrease of the fb NADH (shifted toward the blue Fig.2B and phasor plots representation Fig. 2C) in cells plated on activating antibodies (Fig. 2B, lower panel). These results were confirmed in primary pre-activated human helper T cells (Fig. S3). Altogether these data suggest that anti-CD3+CD28 activation of Jurkat T cells and primary human CD4^+^ T cells induces a rapid shift (from 10 minutes) along the metabolic trajectory from OXPHOS to glycolysis. To confirm that the metabolism induced in the different conditions is reflected by the fb NADH in T cells, we treated the cells with rotenone, which is a drug inhibiting the complex I of the mitochondrial electron transport chain, thus blocking OXPHOS (Fig. S2A). As shown in Fig. S2B-D, rotenone treatment in resting Jurkat T cells significantly decreased the fb NADH, reflecting a shift from OXPHOS to glycolysis in the treated T cells. Conversely, treatment of activated Jurkat T cells with 2-deoxy-d-glucose (2-DG), which interferes with d-glucose metabolism induced a strong increase in fb NADH (Fig. S2B, S2C, S2E) reflecting the inhibition of glycolysis. This was true both in Jurkat (Fig. S2) and primary human CD4^+^ T cells (Fig. S3E).

Altogether, these data show that FLIM analysis of the fb NADH in T cells can reveal dynamic variation of T cell metabolism upon activation.

### Monitoring fraction of bound NADH in single Jurkat cells over time reveals a rapid switch toward glycolysis upon activation

Analysis described in the previous paragraph showed a heterogeneity among Jurkat T cells in the NADH fluorescence lifetimes (Fig. 2) both at resting state and in activating conditions. We thus performed analysis both at the pixel level and single cell level by considering the metabolic signature of the entire cell in 3D and averaging the fb NADH inside the mask defined by the LifeAct-mCherry fluorescence (Fig. S1B). The fb NADH of single cells in each Z was determined (Fig. S1C) by averaging the fb NADH over the segmented region of interest of the single cells in each Z plane. The mean value of fb NADH for the entire cell in 3D was calculated by averaging the fb values over all the Z planes. As shown in Fig. 3A and 3B we could follow the NADH intensity and fb NADH in each single cell longitudinally. Examples of 3 cells in the same field of view in resting (1 to 3) and activating (4 to 6) conditions are represented (Fig. 3C), which show that there was no significant change of the fb NADH in cells plated on PLL whereas there was a progressive decrease of the fb NADH in single cells activated on anti-CD3+CD28 coated surfaces. Analysis of ∼700 cells confirmed that the fb NADH was significantly decreased in activated Jurkat T cells as compared to resting cells at all time points (10, 30 and 60 minutes) (Fig. 3D). It also showed that the fb NADH was significantly decreasing over time in activated T cells. This result was also confirmed in human primary CD4^+^ cells, which showed no significant changes of the fb NADH under resting conditions but a decrease under activating conditions at all time points (Fig. S3C).

**Fig. 3:**
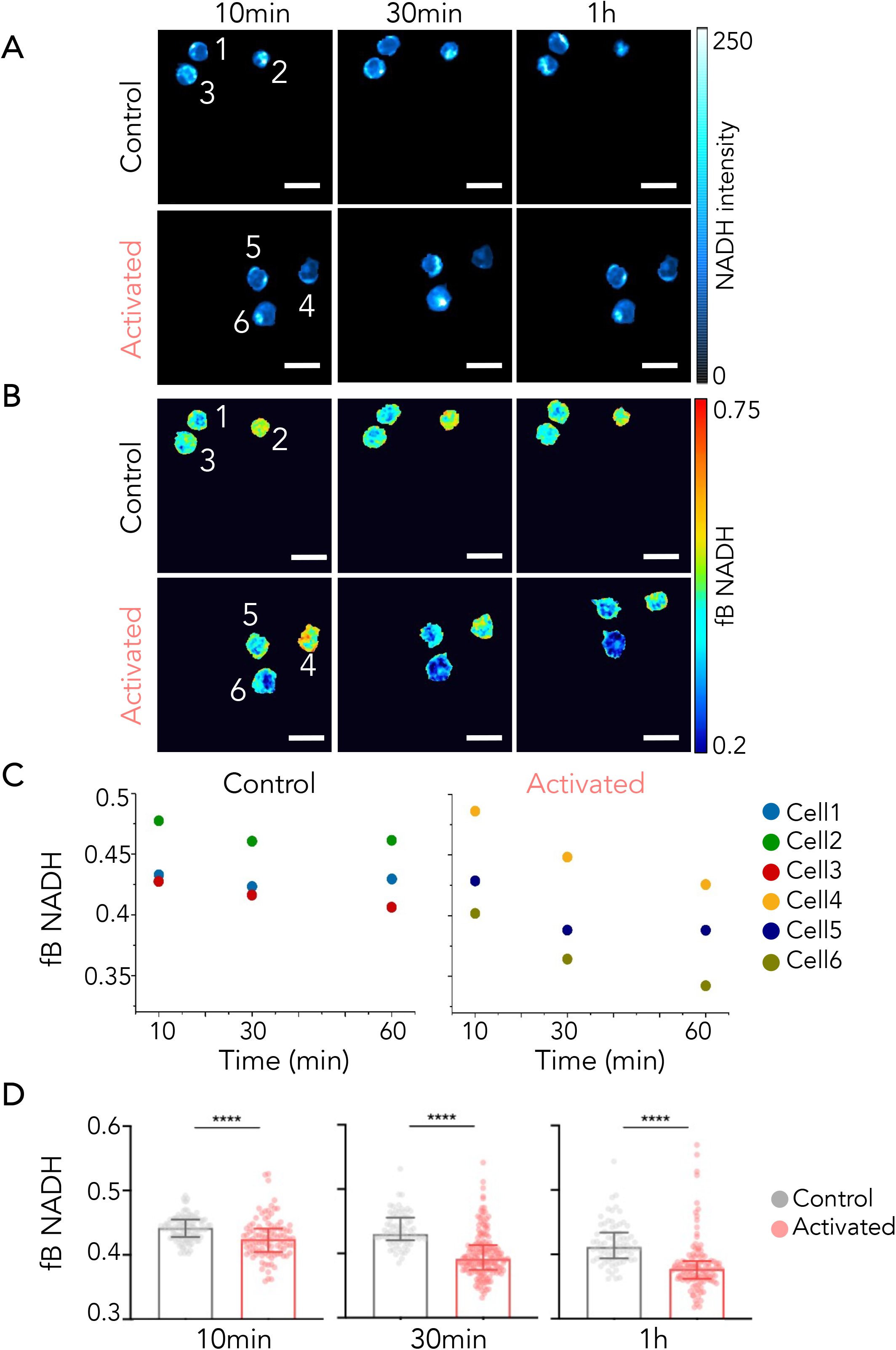
Single cell analysis of NADH FLIM reveals metabolic shift over time during T cell activation. **A**. Representative images of NADH intensity and **B**. maps of fraction of bound NADH of T cells in control and activated condition at three time points (10 min, 30 min and 1 hour). **C**. Fraction of bound NADH of three representative single cells in control and activated conditions over time (10 min, 30 min and 1 hour). **D**. Quantification of fraction of bound NADH over time of control and activated T cells (10 min, 30 min and 1 hour). Data are presented by median with interquartile range. T-test: ****P<0.0001

Altogether, these results suggest that T cell activation induces a rapid switch along the metabolic trajectory from OXPHOS to glycolysis, in the order of minutes that remains stable at least an hour.

### Three-dimension (3D) analysis of the fraction bound NADH in Jurkat T cells

As described previously (Fig. S1), the analysis of the fb NADH was performed both in time and space, i.e. at different Z planes of single T cells. We studied the 3D distribution of the intracellular fb NADH in resting and activated conditions at the 3 chosen time points. As shown in representative images (Fig. 4A) and analysis of the intracellular fb NADH distribution in 700 Jurkat T cells (Fig. 4B), the fb NADH distribution showed a characteristic distribution in both quiescent and activated T cells. In both cases, the two highest values of fb NADH were found at the bottom plane (the contact zone with the slide: 0μm) and top of the cells that mainly contain mitochondria, while the central zone, where the nucleus is predominant, presents the lowest values of fb NADH (Fig. 4B). Moreover, for most Z planes, the fb NADH in activated T cells was lower than the fb NADH in resting cells (Fig. 4B), confirming the results obtained by averaging the fb values of all Z planes (Fig. 3) and suggesting no specific cellular compartmentalization of the activity. The difference between fb NADH in resting and activated T cells was accentuated over time (Fig. 4B) in every location of the cell, showing that the shift toward a glycolytic activity that starts few minutes after CD3/CD28 activation is steady. The distribution of fb NADH in the activated cells (Fig. 4C) shows asymmetry between the cell side in contact with the slide (the surrogate immune synapse) and its opposite, with a fb NADH that is higher in the contact zone than in the top of the cells. This asymmetry suggests that there is relatively more OXPHOS at the synaptic side than in the rest of the cell, whereas in resting cells the distribution of fb NADH is symmetric (Fig. 4C). This might be due to the polarized distribution of mitochondria at the immune synapse^35–37^.

**Fig. 4:**
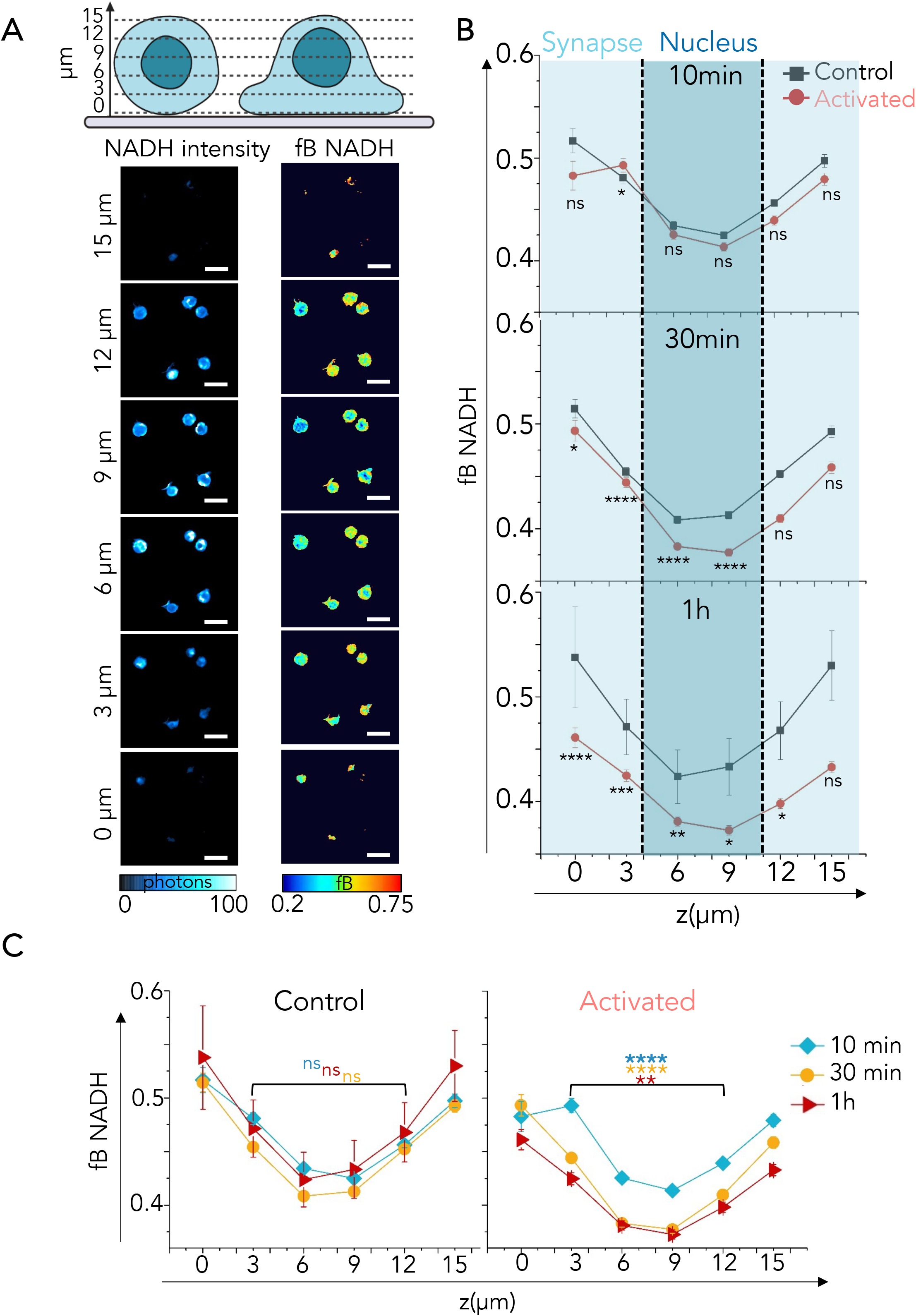
Three-dimensional metabolic pattern in T cells during activation revealed by NADH FLIM. **A**. Representative images of NADH intensity and fraction of bound NADH of single T cell in control at 10 mins in different Z planes. **B-C**. Quantification of fraction of bound NADH in control and activated condition in every Z plane of the cell over time (10 min, 30 min and 1 hour). Data are presented by mean ±SEM. T-test: *P ≤ 0.05, **P ≤ 0.01, *** ≤ 0.001, **** ≤ 0.0001. **B**. Statistics are indicated for each Z plane between control and activated conditions. **C**. Statistics are indicated for each timepoint between Z = 3μm and Z = 12μm.

Altogether, these data show that 2P-FLIM of NADH in T cells can reveal temporal and spatial dynamic changes in their metabolism at a single cell level and in subcellular compartments.

### Following T cell spreading at the immune synapse together with the fraction bound NADH

To better follow the shape of the T cells during activation and define the cell binary “mask” described in Fig. S1B, we transduced Jurkat T cells with LifeAct-mCherry that labels filamentous actin. Within minutes of contact with an activating surface, the actin networks are dynamically and drastically remodeled^7^ forming a bright F-actin ring at the cell periphery (Fig. 5A).

**Fig. 5:**
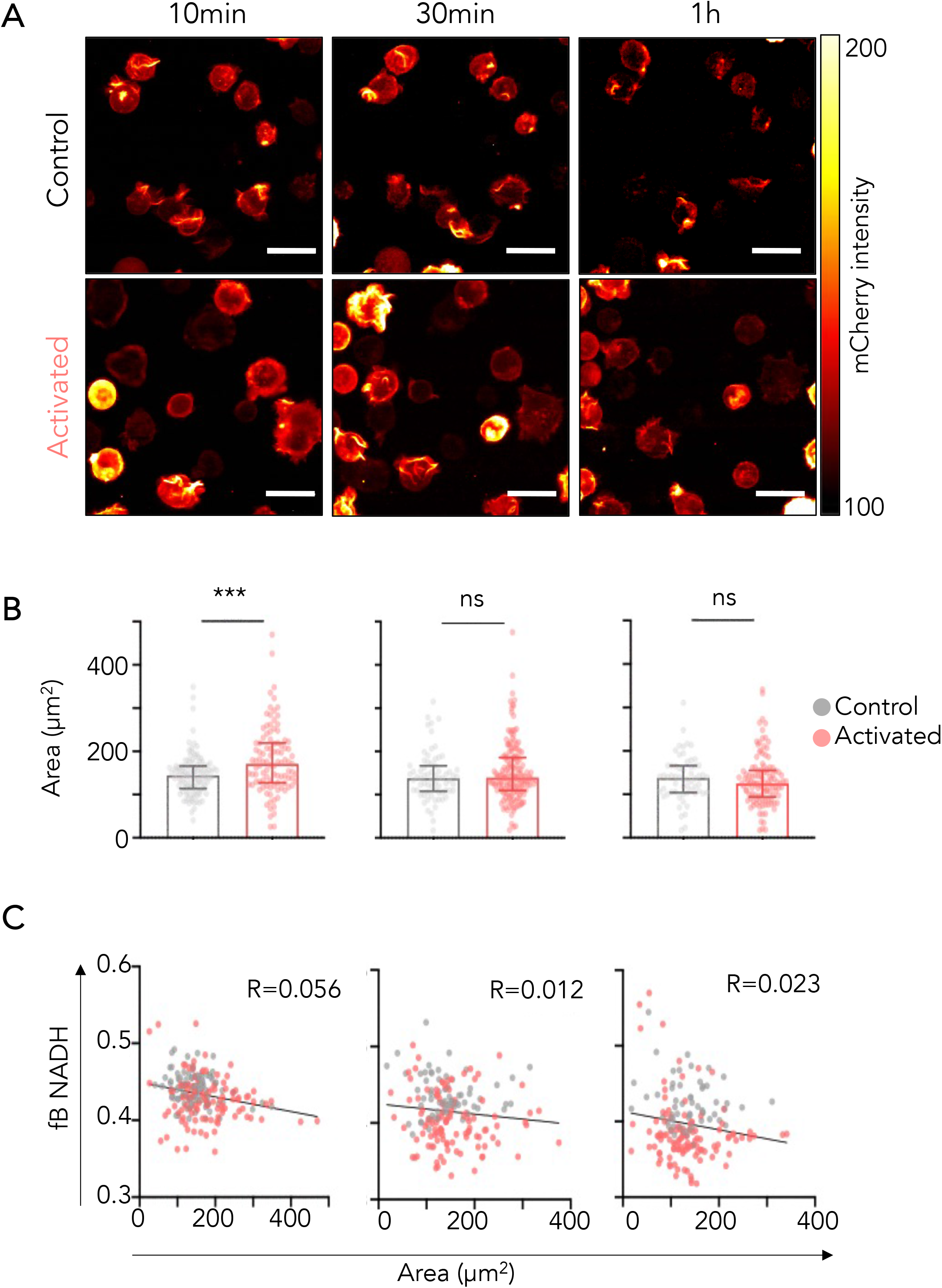
T cell surface spreading during activation. **A**. Representative images of LifeAct-mCherry intensity of T cells in control and activated condition at three time points (10 min, 30 min and 1 hour). **B**. Quantification of T cell area over time (10 min, 30 min and 1 hour) of control and activated T cells. Data is presented by median with interquartile range. **C**. Correlation scatter plot between areas of LifeAct-mCherry and fraction of bound NADH of control and activated T cells over time (10 min, 30 min and 1 hour). T-test: *** P ≤ 0.001

We thus simultaneously measured in 3D the spreading of T lymphocytes on the control (PLL) or activating surface (anti-CD3+anti-CD28) and of the fb NADH. As shown in representative images in Fig. 5A and quantification in Fig. 5B, activation of Jurkat T cells induced a spreading of T cells on the activating surface at 10 minutes. At later time points, cells retracted as previously described^38^.To investigate the relationships that may exist between actin remodeling at the immune synapse and changes in T cell metabolism, we plotted fb NADH as a function of cell area for all the cells (control and activated conditions) at the 3 time points. As shown in Fig. 5C, our data show that there is a negative correlation between cell spreading and fb NADH, which suggests that cells that remodel their actin cytoskeleton the most actively are more glycolytic. This is more striking at 10 min, when cells are spreading the most. We reproduced these results in human primary CD4^+^ T cells (Fig. S3D) measuring the area of the cells using the NADH intensity since the cells did not express LifeAct-mCherry.

Altogether, these results show that two-photon fluorescence lifetime imaging (FLIM) of NADH in T cells can reveal dynamic changes in their metabolism at a single cell level and can be combined with the analysis of other parameters, such as actin remodeling, that can be followed by fluorescence imaging.

## DISCUSSION

Metabolic responses in T cells are highly dynamic and heterogeneous both temporally and spatially, which make them difficult to analyze. Therefore, quantitative, high-resolution, label-free methods to noninvasively examine metabolic processes in cells are particularly needed to better characterize and elucidate the role of different metabolic pathways in normal and pathophysiological conditions. Two-photon excited fluorescence is well appropriated for sensitive, quantitative, label-free, high-resolution assessments of metabolic activity *in vitro* and *in vivo*^17,39,40^. Several endogenous molecules, including NADH are autofluorescent. The fluorescence lifetime of NADH varies whether it is free or bound, thus can be used as a metabolic indicator^18^. NADH in the mitochondria is known to be primarily protein bound, resulting in a fraction bound NADH that has a longer lifetime than the free NADH in the cytosol and nucleus ^41,42^. Thus, changes in the lifetime of NADH has been attributed to alterations in the relative levels of oxidative phosphorylation and glycolysis ^14,19,43^. In this study, we implemented Two-Photon excitation of NADH and FLIM to map the temporal and spatial metabolic patterns in isolated human T cells, i.e., the widely used Jurkat cell line ^44^ and human primary CD4^+^ T cells to follow the metabolic changes upon activation. Using metabolic inhibitors (Fig. S2 and S3), we determined the specificity of the fb NADH measurement to defined metabolic pathways: rotenone which induces a shift from OXPHOS to glycolysis by blocking the respiratory chain increased fb NADH in quiescent cells. Besides, 2-deoxy-d-glucose (2-DG), which interferes with glucose metabolism induced a decrease in fb NADH in activated T cells that revealed the shift toward more OXPHOS. Our results show that T lymphocyte activation by the TCR/CD3 and the CD28 molecules rapidly (in 10 minutes) induces a decrease in fb NADH, witnessing a metabolic shift toward more glycolysis. The measurement of fb NADH in live cells enables an early observation of the metabolic switch upon T cell activation while most of the studies analyze metabolic changes after 24h to 48h of activation. The metabolic shift toward glycolysis reported in these cells can thus be due to transcription of genes encoding enzymes controlling glycolysis. Indeed, we have shown that human T lymphocytes activated by anti-CD3 and anti-CD28 activating antibodies increase transcription of genes encoding: Phosphofructokinase, Glyceraldehyde-3-Phosphate Dehydrogenase, Phosphoglycerate Kinase 1, phosphoglucomutase, Enolase and Pyruvate kinase ^45^, which are all enzymes controlling glycolysis. In the present study, the rapidity of the metabolic shift toward glycolysis strongly suggests that the process is transcription independent. Our results, which show a negative correlation between cell spreading and fb NADH, suggest a causal effect between actin remodeling and glycolytic activity of T cells. Indeed, direct interactions between glycolytic enzymes and filamentous actin have been reported and actin remodeling shown to regulate these enzymatic activities (reviewed in ^46^). Such a regulation is exemplified by a study reporting that insulin induces the release of enolase A from F-actin via a PI3K and Rac dependent actin remodeling. Free aldolase then leads to an increase in total aldolase activity, driving glycolytic flux in insulin activated cells^47^. Interestingly, TCR/CD3 and CD28 activation also induce Rac and PI3K activation in T lymphocytes ^48–50^. It would thus be interesting to explore if the rapid cytoskeleton remodeling observed in activated T cells also controls the availability of glycolytic enzymes in these cells. Alternatively, there might be no causal effect: spreading of T cells and increase in F-actin would reflect T cell activation: the more activated the cell, the more glycolytic it is.

This method was already used to show that the autofluorescence decay of NADH is sensitive to T cell subtypes and activation state^12^. Heterogeneity of NADH lifetime was reported in T lymphocytes in a given donor and among donors, it was also different between CD3^+^CD4^+^ and CD3^+^CD8^+^ activated T cells ^12^. Moreover, the fraction of free NADH was lower in quiescent T cells from the blood of healthy donors than in T cells activated for 24 to 72h with tetrameric antibodies (anti-CD2/CD3/CD28) ^12^ and the authors of the study reported that the lifetime of NADH was consistently the most important biomarker for differentiating quiescent and activated T cells ^12^. Our results showing heterogeneity in the fb NADH both in Jurkat cells, which are known to present clonal heterogeneity ^51^ and in primary activated CD4^+^ T cells are in accordance with this published study. The decreased fb NADH we observed in activated T cells is also consistent with this published study although the timing was different. One important difference is that we only measured the lifetime of NADH focusing on the OXPHOS and glycolysis pathways, whereas in the cited study ^12^, the Walsh at al. measured multiple metabolic intrinsic biomarkers (lifetime of NADH, lifetime of flavin adenine dinucleotide (FAD) and optical redox ratio (defined as NADH/(NADH+FAD)) that provide complementary information about different metabolic pathways and phenotypes including fatty acid synthesis and glutaminolysis ^52,53^. Moreover, we concentrated on the early metabolic changes. Using fluorescence imaging of both endogeneous NADH and FAD, the authors reported an initial (at 30 minutes) decrease of the optical redox ratio in T lymphocytes that then increases over time (from 2h to 24h) ^12^. It is thus difficult to directly compare the results we report herein with the results of Walsh et al due to different T cell activation, imaging timing and biomarker measurements. Yet, it would be of high interest to characterize and compare the spatial dynamics of metabolism switch in T cells induced, at early time points, by different activators (coated, soluble crosslinked antibodies, tetramers), using multiparametric metabolic imaging with Two-photon Fluorescence Lifetime Microscopy (2P-FLIM) of the coenzymes NADH and FAD simultaneously ^22^. Of note, the rapid decrease in fb NADH in activated T cells that reflects an increase in the glycolytic activity of these T lymphocytes was reported in another model. In this case, the authors used the Seahorse Extracellular Flux Analyzer, and demonstrated, in mouse T lymphocytes, a rapid (within minutes) increase of glycolysis upon activation with crosslinked anti-CD3 and anti-CD28 antibodies^11^. This rapid aerobic glycolysis was transcription independent and due to activation of pyruvate dehydrogenase kinase 1 (PDHK1).

One other interest of the method and work we present is that it allows a spatial visualization of the metabolism inside single cells. In our study, we show that there is a gradient of fb NADH in T lymphocytes. In non-activated T lymphocytes, the fb NADH is evenly distributed in the cytoplasm while the central zone, representing mostly the nucleus, displays the lowest values (Fig. 4B). This can be explained by the absence of mitochondria in the nuclei. One interesting aspect of our results is the difference in the distribution of fb NADH in the activated cells, which shows asymmetry between the cell side in contact with the slide and its opposite. The fb NADH is higher at the contact zone than at the top of the cells, whereas in resting cells the distribution of fb NADH is symmetric at the two poles of the cells. This probably reflects the polarization of T lymphocytes toward the activating side, also called immune synapse, which induces rapid polarized transport of mitochondria at the immune synapse ^35,54^. This asymmetrical distribution of fb NADH is more pronounced at early time points (10 and 30 minutes) than at later time point (1h). This might reflect the fact that mitochondria first accumulate in the immediate vicinity (<200nm) of the immune synapse and then follow a retrograde movement toward the distal end of the T cell ^37^. It would be interesting to study if the polarity of T cells affects their metabolism. To do so, we could compare the lifetime of NADH induced by coated antibodies (as herein), which mimics the contact with an antigen presenting cell and induce polarization and morphological changes of the T cells, with data obtained in non-polarizing conditions (soluble antibodies).

T lymphocytes are poised to rapidly respond to diverse stimuli, in different nutrient contexts, and different physiological and pathological conditions. This ability to rapidly adapt to many situations must be accompanied by rapid changes in their cellular metabolism to fulfill their need of energy. It is thus important to develop and implement methods, which can document the metabolism changes in these immune populations. We expect that the use of Two-photon Fluorescence Lifetime Microscopy (2P-FLIM) of the coenzyme NADH will provide valuable information on diverse T cell populations from non-pathological and pathological samples and in response to diverse stimulations.

## Acknowledgements

We would like to thank Xavier Lahaye for providing us with peripheral blood mononuclear cells and the members of the “Integrative analysis of T cell activation” (Institut Curie) for support and discussions. We thank all the members of the “advances microscopies” group at the Laboratory for Optics and Biosciences for helpful discussions and advice. We thank Pierre Mahou and Emmanuel Beaurepaire for scientific discussions and technical help.

## FIGURES LEGENDS

**Fig. S1:**
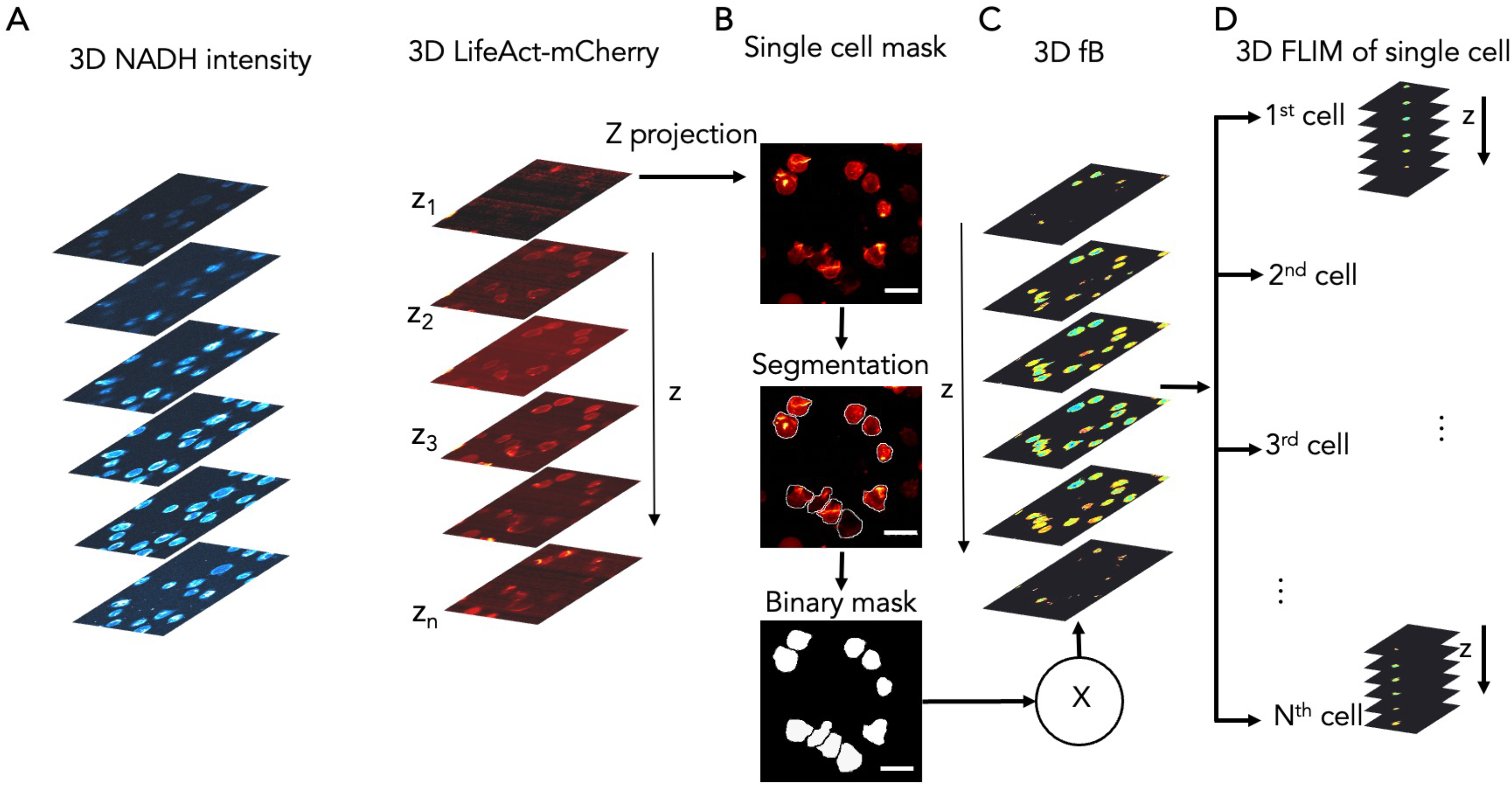
Workflow of 3D FLIM for single cell analysis. **A**. NADH FLIM and LifeAct-mCherry Z-stack image **B**. The z projection is used to create single cell masks. **C**. 3D of the fraction of bound NADH **D**. The single cell mask is applied to the 3D FLIM raw data of fraction of bound NADH (fb NADH) in order to extract the FLIM data of single cell for every Z plane.

**Fig. S2:**
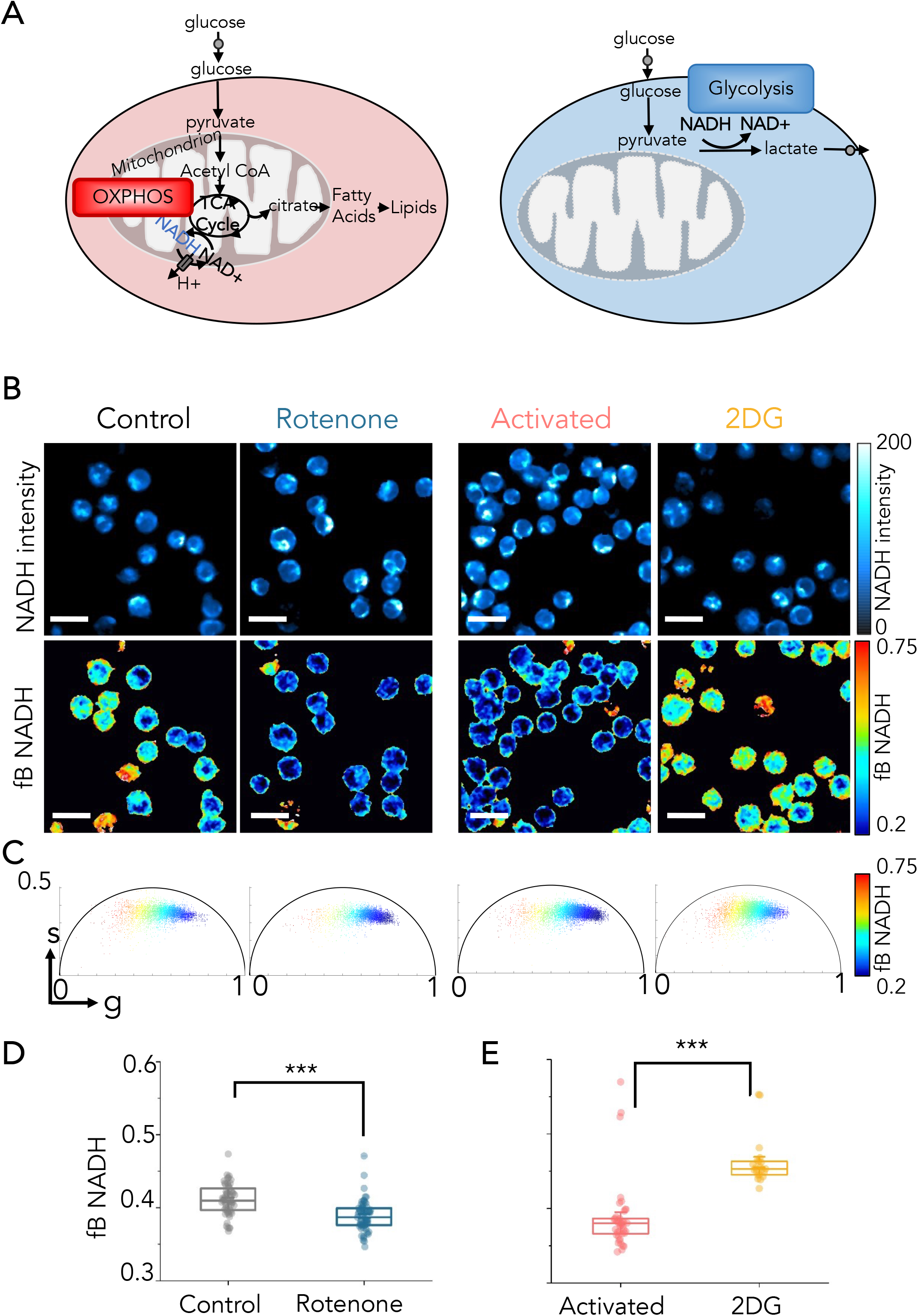
Metabolic trajectory in Jurkat T cells. **A**. Schematic representation of the metabolic pathways inside the cells: oxidative phosphorylation (OXPHOS) and glycolysis. Glucose breakdown through glycolysis and the TCA (tricarboxylic acid) cycle generates reduced NADH and FADH_2_. Cells with a glycolytic phenotype generate ATP rapidly in the cytoplasm and are characterized by a high ratio of NADH/NAD^+^ and a low fraction of bound NADH. Cells relying on oxidative phosphorylation convert glucose to pyruvate, which is then oxidized in the TCA cycle generating the majority of ATP in the mitochondria and are characterized by a low ratio of NADH/NAD^+^ and a high fraction of bound NADH. **B**. Intensity images (top panel) and fraction of bound NADH maps (bottom panel) of Jurkat T cells in control condition and with rotenone as well as in activated condition and with 2-Deoxy-D-glucose (2DG). *Rotenone blocks the respiratory chain* and leads to a decrease fraction of bound NADH, while 2DG prevents glycolysis and leads to an increase of bound NADH. **C**. Representative phasor plot of Jurkat T cells in different conditions: control, control treated with rotenone, activated, activated treated with 2-Deoxy-D-glucose. **D**. Decrease of fraction of bound NADH after rotenone treatment in control. **E**. Increase of fraction of bound NADH in activated Jurkat T cells with 2-Deoxy-D-glucose. Data are presented by mean with SEM. T-test: *** P ≤ 0.001

**Fig. S3.**
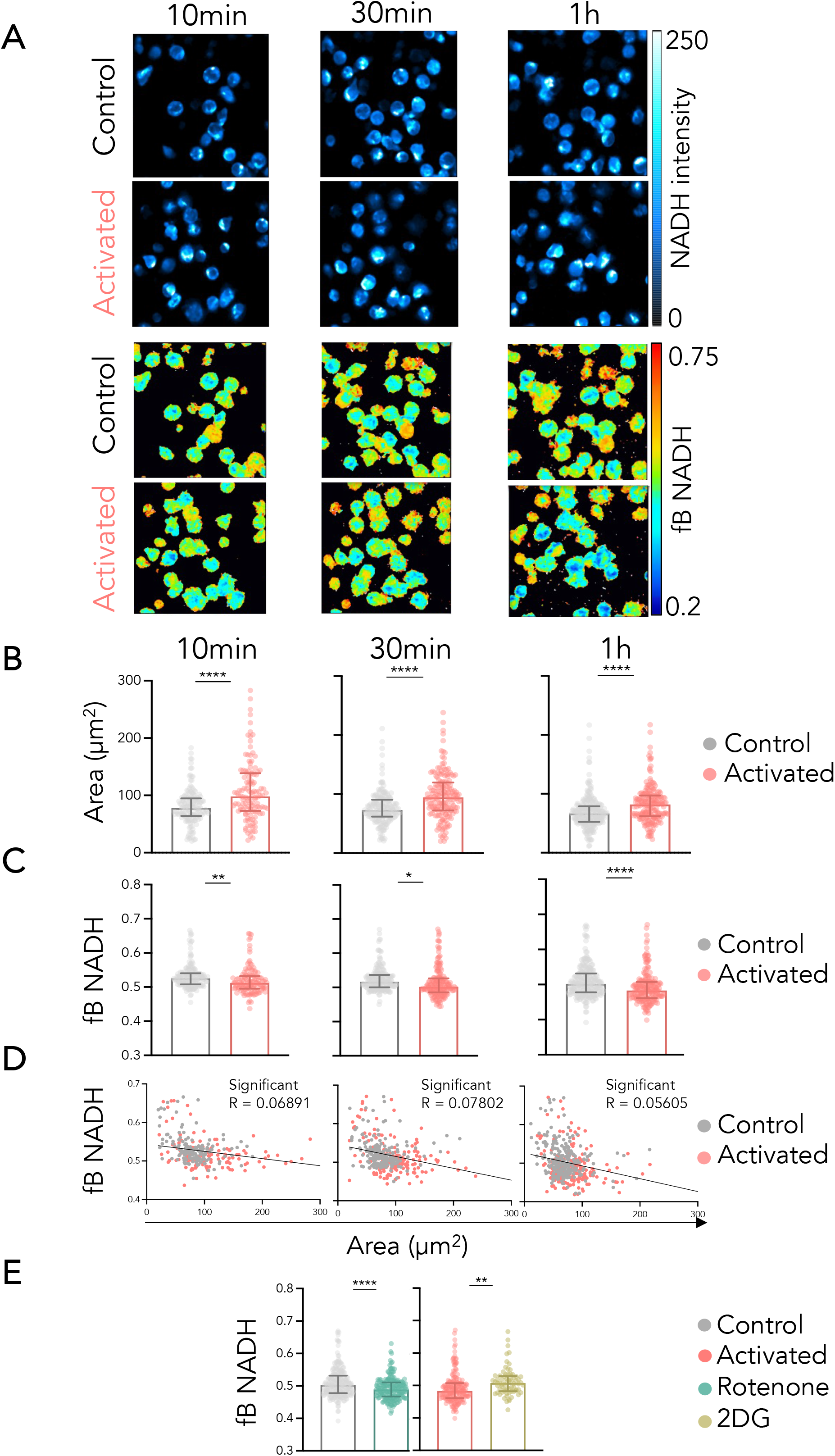
Single cell analysis of NADH FLIM reveals metabolic shift over time during primary T cell activation. **A**. Representative images of NADH intensity (top panel) and maps of fraction of bound NADH images (bottom panel) of primary T cells in control and activated condition at three time points (10 min, 30 min and 1 hour). **B**. Quantification of primary T cell area over time (10 min, 30 min and 1 hour) of control and activated T cell. Data are presented by median with interquartile range. **C**. Quantification of fraction of bound NADH over time of control and activated T cell (10 min, 30 min and 1 hour). Data are presented by median with interquartile range. **D**. Correlation scatter plot between areas of LifeAct-mCherry and fraction of bound NADH of control and activated T cells over time (10 min, 30 min and 1 hour). **E**. Decrease of fraction of bound NADH after rotenone treatment in control and increase of fraction of bound NADH in activated cells with 2-Deoxy-D-glucose. Data are presented by median with interquartile range. T-test: *P ≤ 0.05, **P ≤ 0.01, ****P≤ 0.0001. N = 1 healthy donor run in duplicates.

